# Reduced Achilles tendon stiffness in aging persists at matched activations and associates with higher metabolic cost of walking

**DOI:** 10.1101/2023.11.27.568808

**Authors:** Jason R. Franz, Rebecca L. Krupenevich, Aubrey J. Gray, John A. Batsis, Gregory S. Sawicki

## Abstract

The mechanisms responsible for increased walking metabolic cost among older adults are poorly understood. We recently proposed a theoretical premise by which age-related reductions in Achilles tendon stiffness (k_AT_) can disrupt the neuromechanics of calf muscle behavior and contribute to faster rates of oxygen consumption during walking. The purpose of this study was to objectively evaluate this premise. We quantified k_AT_ at a range of matched activations prescribed using electromyographic biofeedback and walking metabolic cost in a group of 15 younger (age: 23±4 yrs) and 15 older adults (age: 72±5 yrs). Older adults averaged 44% less k_AT_ than younger adults at matched triceps surae activations (p=0.046). This effect appeared to arise not only from altered tendon length-tension relations with age, but also from differences in the operating region of those length-tension relations between younger and older adults. Older adults also walked with a 17% higher net metabolic power than younger adults (p=0.017). In addition, we discovered empirical evidence that lesser k_AT_ exacts a metabolic penalty and was positively correlated with higher net metabolic power during walking (r=-0.365, p=0.048). These results pave the way for interventions focused on restoring ankle muscle-tendon unit structural stiffness to improve walking energetics in aging.

## Introduction

When walking at the same speed, older adults (≥65 years) consume metabolic energy much faster than younger adults (e.g., 18-35 years), which can accelerate fatigue and reduce independence and quality of life)1-5. Unfortunately, the mechanisms responsible for the increasing metabolic cost in the aging process are poorly understood. Given their role in powering locomotion, muscle-tendon units (MTUs) spanning the ankle have been shown through experimental and computational modeling studies to play a role in governing metabolic cost^6-10^. In particular, neuromechanical interaction between triceps surae muscle function and Achilles tendon stiffness is critical for tuning positive power generation during push-off which is in turn responsible for as much as much as half of the cost of walking^11,12^. This is significant, as most human subject comparisons have shown *in vivo* evidence of reduced Achilles tendon stiffness (k_AT_) with age^13-17^. Indeed, we recently proposed a theoretical mechanistic pathway by which age-related reductions in Achilles tendon stiffness can disrupt the neuromechanics of calf muscle behavior and contribute to faster rates of oxygen consumption during walking^17^. This viewpoint would pave the way for interventions focused on restoring ankle MTU structural stiffness to improve walking energetics in aging. However, no study to date has moved beyond a theoretical proposal to provide direct empirical data in support of this association.

Basic physiological principles dictate that a muscle operating at shorter than optimal lengths will require greater excitations to generate a given amount of force. Conceptually, age-related reductions in k_AT_ would elicit more tendon stretch for a given load. Older and younger adults walk with similar ankle joint kinematics and thus similar triceps surae muscle-tendon unit lengths^18^. Thus, for a given activation, the more compliant Achilles tendon of older adults would compel shorter triceps surae muscle operating lengths. These predictions of shorter operating lengths are supported by computational simulations and experimental comparisons between younger and older adults^7,10,19,20^. Ultimately, it is this neuromechanical cascade, arising from reduced k_AT_, that we have proposed contributes at least in part to a higher metabolic cost of walking for older adults.

Net ankle joint function is thought to arise from a combination of activation-independent (i.e., Achilles tendon) and activation-dependent components (i.e., muscle). Previous work from our lab indicates that ankle MTU stiffness is regulated via activation-dependent changes in triceps surae length-tension behavior^21,22^. Tendon length-tension relations are non-linear at lower tissue strains and the effective stiffness “seen” by muscle can thus vary as a function of activation. Given the complexity of age-related differences in ankle MTU neuromechanics, it is possible that prior studies may have inadvertently compared k_AT_ between younger and older adults at different regions of their respective force-length curves. However, it is unknown how aging effects on Achilles tendon stiffness varies as a function of triceps surae activation. Understanding the landscape of these age-related differences is an important prerequisite to understanding their influence on walking energetics.

Therefore, the purpose of this study was two-fold. First, we aimed to quantify age-related differences in apparent Achilles tendon stiffness across a range of matched muscle activations using electromyographic biofeedback. Second, we aimed to determine the relation between Achilles tendon stiffness at matched activations and the metabolic cost of walking across age. We hypothesized that: (1) independent of age, Achilles tendon stiffness would increase with higher triceps surae activations, consistent with a shift from the nonlinear to linear region of the length tension curve, (2) older adults would exhibit lesser Achilles tendon stiffness compared to young adults at matched activations, and (3) lesser Achilles tendon stiffness would positively correlate with a higher metabolic cost of walking.

## Methods

### Participants and study design

We recruited 15 younger adults (23±4 years, 8F/7M, 72.9±14.2 kg, 1.7±0.1 m) and 15 older adults (age: 72±5 years, 9F/6M, mass: 74.6±17.1 kg, height: 1.69±0.10 m) who participated after providing written, informed consent according to the University of North Carolina Biomedical Sciences Institutional Review Board. Prior to participation, subjects were screened and excluded if they reported neurological disorders affecting the lower-extremity, musculoskeletal injury to the lower-extremity within the previous six months or were currently taking medications that cause dizziness. Two experimental sessions were performed: one for isolated contractions and the determination of Achilles tendon stiffness, and the second for determining net metabolic power during treadmill walking (**Fig. 1**). For all isometric and isokinetic tasks, participants were seated in a dynamometer (Biodex, Shirley, New York) at 85° hip flexion and 20° knee flexion. For the walking task, participants walked on a Bertec split-belt treadmill at a fixed speed of 1.25 m/s.

**Figure 1.**
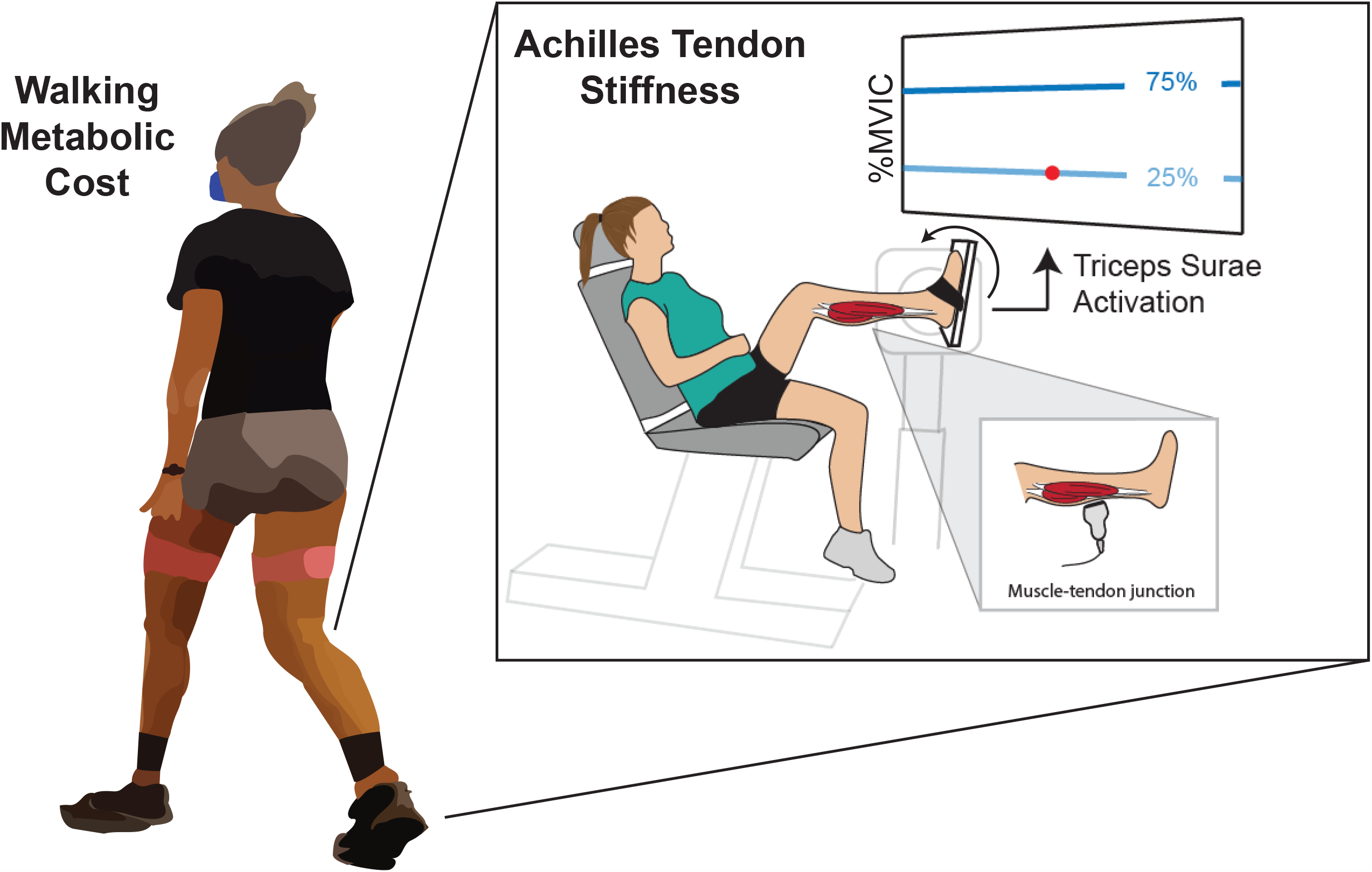
Methodological diagram showing the primary experimental paradigms and outcome measured used in this study. Subjects performed isokinetic ankle dorsiflexion tasks while responding to electromyographic (EMG) biofeedback designed to matched different activation levels between younger and older participants. Motion capture-guided ultrasound imaging and dynamometry were combined to estimated Achilles tendon stiffness. We correlated those stiffness values to net metabolic power estimated from treadmill walking at a fixed speed of 1.25 m/s.

### Isolated contractions: protocol and measurements

We placed surface recording electrodes (Delsys, Natick, Massachusetts) on participants’ right soleus and lateral gastrocnemius after prepping the site by shaving and abrading the skin. Sensors were secured to the skin using adhesive tape. For the electromyographic (EMG) biofeedback described below, soleus and lateral gastrocnemius data sampled at 1000 Hz were demeaned, full-wave rectified and bandpass filtered (20-450 Hz). Soleus and lateral gastrocnemius data were averaged and defined as “triceps surae activation” throughout the rest of the manuscript. Simultaneously, we recorded the 3D trajectories of 20 retroreflective markers placed on the right leg and clusters affixed to each of two ultrasound transducers using motion capture operating at 100 Hz (Motion Analysis, Santa Rose, CA). Participants first donned one 60-mm linear array transducer (LV7.5/60/128Z-2, Telemed Echo Blaster 128, Lithuania) secured to the right shank via a custom 3D-printed housing and positioned to image the medial gastrocnemius muscle-tendon junction (MTJ) at 61 frames per second. Participants also donned a 38-mm transducer (L14-5W/38; Ultrasonix corp, Richmond, British Columbia), operating at 70 frames per second, placed on the Achilles free tendon distal to the soleus muscle-tendon junction. Biodex position and torque data were recorded at 1000 Hz in synchrony with motion capture data. A voltage sync signal from each ultrasound machine identified the start and stop of their acquisitions.

Participants completed a series of plantarflexor maximum voluntary isometric contractions (MVICs) at six ankle dorsiflexion (DF) and plantarflexion (PF) joint angles (20° DF, 10° DF, 0°, 10° PF, 20° PF, 30° PF). If a participants’ maximal dorsiflexion range of motion (ROM) was greater than 25°, a seventh joint angle was added that was equal to their maximum dorsiflexion ROM. We corrected the net ankle joint moment for gravitational and passive moments. Data from isometric contractions were used to: (1) determine the coefficients for subject-specific moment arm equations (all joint angles), and (2) establish a reference maximum triceps surae activation for the isokinetic EMG biofeedback trials to follow (neutral joint angle).

Thereafter, we displayed triceps surae activation on a monitor positioned in front of the participant in real-time with a target line corresponding to a percent of MVIC activation (**Fig. 1**). Participants performed eccentric isokinetic plantarflexor contractions at 20 deg/s for two activation conditions: 25% MVIC and 75% MVIC. Participants were instructed to match their instantaneous EMG (displayed as a dot on the screen) to a target line (a horizontal line across the screen). A third condition was performed during which the dynamometer moved the ankle joint through the prescribed range of motion without active subject resistance (referred to herein as passive rotation). Two trials were collected for each movement. Participants rested between trials and were allowed to practice the movement prior to data collection. All three of these conditions were randomized.

### Isolated contractions: data processing and analysis

We located the position of the medial gastrocnemius MTJ in each frame of ultrasound video using an automated deep-learning algorithm^23^. We co-registered these positions with the position of the calcaneal marker as a surrogate for Achilles tendon insertion, and calculated the Achilles tendon elongation relative to the start of each isokinetic contraction or passive rotation.

The Achilles free tendon ultrasound data and the corresponding motion capture data were then used to estimate subject-specific moment arms. For each ankle joint angle used during isometric testing, we estimated the AT moment arm using previously published procedures as the average perpendicular distance between the tendon’s line of action, registered manually in the ultrasound images, and the transmalleolar midpoint^24^. We then curve-fit those points to calculate AT moment arm as a function of ankle joint angle for each participant.

We estimated time series of AT force by dividing net ankle joint moments measured from the dynamometer by subject-specific AT moment arm. Finally, we calculated AT stiffness (i.e., k_AT_) as the linear slope of the relation between AT force and AT elongation between 20 and 80% of each participants’ dorsiflexion range of motion.

### Treadmill walking: net metabolic power

During a 5-min standing trial and a 5-min habitual walking trial, we sampled breath-by-breath rates of oxygen consumption and carbon dioxide production using a COSMED K5 indirect calorimetry system (COSMED, Rome, Italy). For each, we computed the average rates of gas exchange over the final two minutes of each collection. We used standard regression equations to estimate whole-body net metabolic power from these average rates of oxygen consumption and carbon dioxide production^25^. Finally, we subtracted standing values from walking metabolic power to calculate net metabolic power, which we normalized to participant body mass (W/kg).

### Statistical Analysis

The primary outcome measures in this study were Achilles tendon stiffness, measured during passive rotation and during eccentric dorsiflexion tasks performed at 25% and 75% MVIC activation, and net metabolic power during walking. We assessed between-group differences in net metabolic power using an independent samples t-test. For k_AT_, we performed a two-way mixed-effects ANOVA to test for a significant effect of age (between-subject effect; old vs. young) and activation (within-subjects factor; passive, 25%, 75%). Partial eta squared (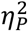) provided the magnitude of the interaction and main effect sizes. We performed all analyses in SPSS and defined significance using an alpha level of 0.05 for all comparisons.

## Results

We did not find a significant interaction between triceps surae activation level and age for k_AT_ (F2,56=1.41, p=0.250, 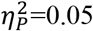) (**Fig. 2**). However, a significant main effect of age (F_1,28_=4.34, p=0.046, 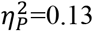) revealed that older adults exhibited, on average, 44% lesser k_AT_ than younger adults across activation conditions. In addition, a significant main effect of activation level (F_2,56_=17.80, p<0.001, 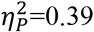) revealed that, across age groups, k_AT_ increased roughly four-fold from passive rotation to 25% MVIC activation, and again by +65% from 25% to 75% MVIC activation. We also found that older adults walked with 17% higher net metabolic power than younger adults (older vs. younger: 3.11±0.36 W/kg vs. 2.66±0.58 W/kg) (p=0.017). We found no correlations between k_AT_ during passive rotation or at 25% MVIC activation and net metabolic power during walking (r≤0.07, p≥0.05) (**Fig. 3A-B**). Conversely, we found that k_AT_ measured at 75% MVIC activation significantly and positively correlated with net metabolic power during walking (r=-0.365, p=0.048) (**Fig. 3C**).

**Figure 2.**
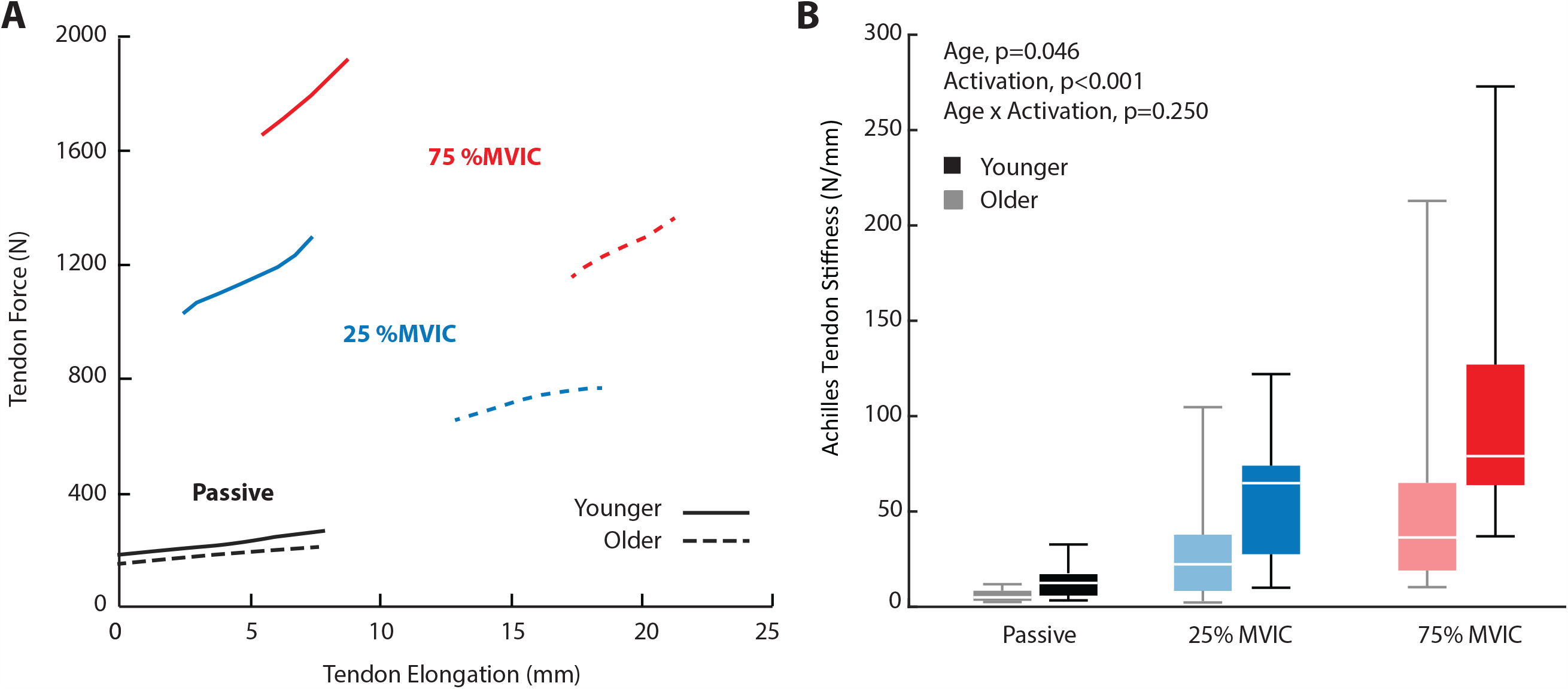
Box-and-whisker plots showing Achilles tendon stiffness (N/mm) derived during isokinetic ankle dorsiflexion performed at rest (i.e., Passive) and at 25% and 75% of peak activation during a maximum voluntary isometric contraction (MVIC).

**Figure 3.**
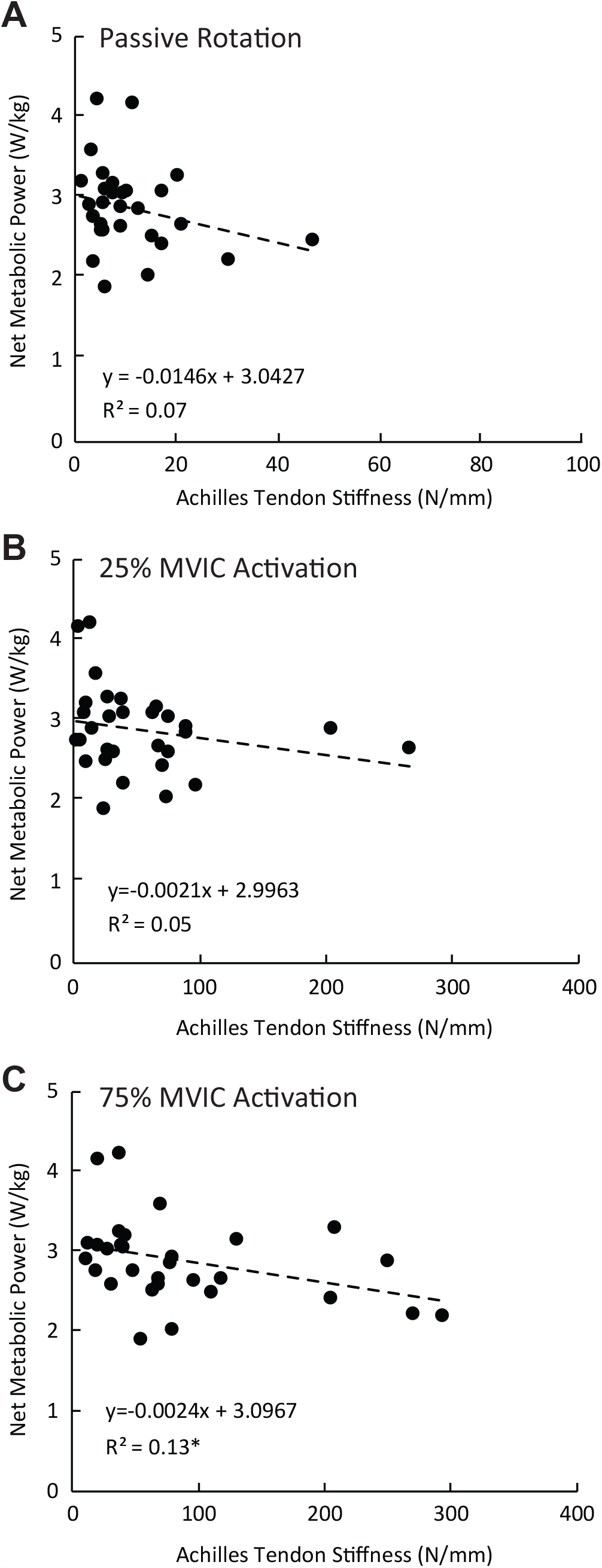
Correlations between Achilles tendon stiffness and net metabolic power during treadmill walking. Values shown represent Achilles tendon stiffness values derived during isokinetic ankle dorsiflexion performed at (A) rest (i.e., Passive) and at (B) 25% and (C) 75% of peak activation during a maximum voluntary isometric contraction (MVIC). Linear fit and statistical outcomes shown for the combined cohort of older and younger adults. Asterisk (*) represents a correlation deemed statistically significant (i.e., p<0.05).

## Discussion

The purposes of this study were to: (1) quantify age-related differences in apparent Achilles tendon stiffness across a range of matched muscle activations using electromyographic biofeedback, and (2) determine the relation between Achilles tendon stiffness at matched activations and the metabolic cost of walking across age. Consistent with our first hypothesis, younger and older adults exhibited Achilles tendon stiffness that increased with increasing triceps surae activation. This outcome reflects the nonlinear length-tension relation for tendon and demonstrates how the mechanics of a seemingly passive elastic tissue can depend on muscle activation. Consistent with our second hypothesis, older adults exhibited, on average, 44% less Achilles tendon stiffness than younger adults at matched triceps surae activations. As we describe in more detail below, we offer an interpretation of these data to implicate not only altered tendon length-tension relations with age, but also differences in the operating region of those length-tension relations between younger and older adults (**Fig. 4**). Finally, consistent with our third hypothesis and as the major contribution of this paper, we discovered empirical evidence that lesser Achilles tendon stiffness exacts a metabolic penalty and is positively correlated with higher net metabolic power during walking.

**Figure 4.**
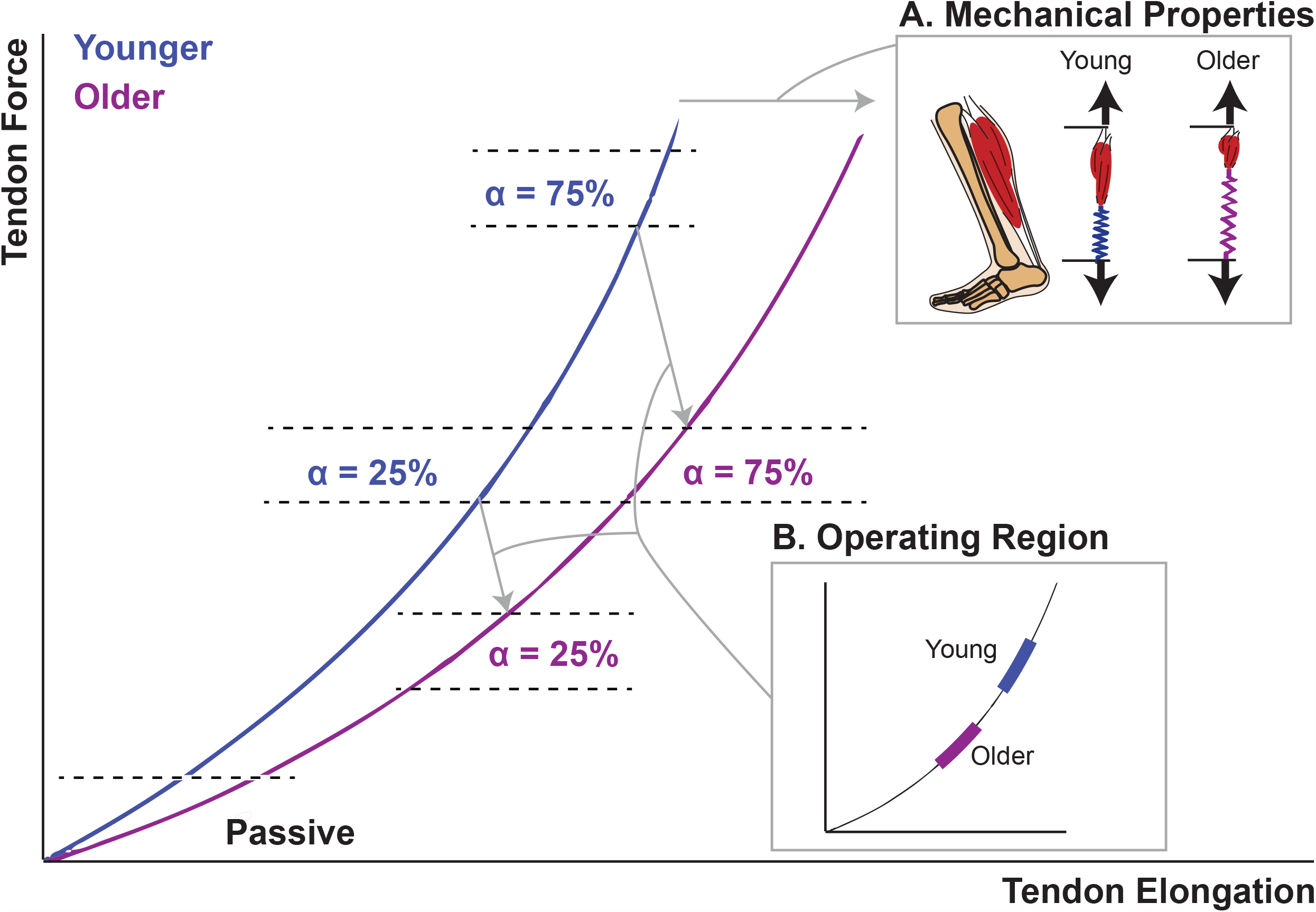
We offer here a conceptual interpretation of our stiffness data from Figure 2 which implicates not only (A) altered tendon length-tension relations with age, but also (B) differences in the operating region of those length-tension relations between younger and older adults.

Our results agree well with previously-published literature. Our controlled-loading experiments generated activation-dependent effects on Achilles tendon stiffness that are highly consistent with those published previously for ankle joint quasi-stiffness in younger and older adults^21^. For example, we showed that ankle joint quasi-stiffness increased by a factor of 6- to 10-fold from passive rotation to isokinetic dorsiflexion at 75% MVIC activation, whereas here k_AT_ increased herein by a factor of 8-9 over the same range of activations. In addition, our average between-group difference in Achilles tendon stiffness (i.e., -44% in older vs. younger adults) is consistent with the majority of *in vivo* human studies (e.g., ^17^). Though, we acknowledge that animal studies sometimes arrive at different conclusions. Finally, our between-group differences in walking net metabolic power (i.e., +17% in older vs. younger adults) agree well with the range of previously published values, which typically average +15-20% (e.g., ^2^)

There are at least two explanations for the observed age-related reduction in Achilles tendon stiffness across the range of matched activations prescribed using EMG biofeedback. **Figure 4** illustrates two likely candidates. The rightward shift evident in our overall data (i.e., indicating longer tendon lengthening per unit increase in force) points to a reduction in k_AT_ as is classically defined by a shallower length-tension curve across the range of activations. We contend that this is typically how age-related reductions in Achilles tendon stiffness are interpreted in the available human literature. Indeed, we have used this interpretation as a mechanism for shorter triceps surae fascicle lengths in older adults during walking^26^, and as evidence for a “structural bottleneck” that may have metabolic implications^17^. Our data support these interpretations and implications. However, our results at matched activations also point to an additional effect of aging; specifically, that a given relative activation, older adults may operate at a shallower region of their tendon length-tension relations than younger adults. Indeed, older adults walk with a diminished push-off characterized by lesser peak ankle moments than younger adults and thus presumably smaller Achilles tendon forces^27,28^. We would encourage follow-on studies to explore experiments in which biofeedback is used to matched both muscle activations and tendon forces between younger and older participants. Such an effort would allow us to more completely map the mechanical demands of everyday walking tasks in older adults to age-related differences in the functional operating behavior of the Achilles tendons.

We have previously demonstrated that: (1) older and younger adults can modulate ankle joint quasi-stiffness via activation-dependent changes in triceps surae muscle length-tension behavior, and (2) despite age-related reductions in ankle joint quasi-stiffness during controlled contractions, older adults maintain quasi-stiffness during via a local adaptive response of increasing triceps surae activation^21^. This discovery motivated the need to deploy EMG biofeedback to match activation in comparisons of Achilles tendon stiffness between older and younger adults. However, our theoretical model in that previous paper purported that, independent of age, ankle joint quasi-stiffness arises from activation-dependent (muscle) and activation-independent (tendon) components. Our results here reveal that the theoretical model motivating that earlier paper was incorrect and neglected consideration that the non-linear portion of the tendon’s length-tension relation is relevant across a range of muscle activations. These results should compel efforts to revisit previous comparisons of ankle-joint quasi-stiffness and its neuromechanical origins, to include the veracity of interpretations of between-condition and between-group comparisons.

From a clinical perspective, our results suggest that interventions designed to augment Achilles tendon stiffness may help to reduce walking metabolic cost for older adults. These results provide experimental credibility to similar recommendations made following the use of musculoskeletal simulations to predict the bioenergetic consequences of walking with reduced Achilles tendon stiffness^7,9,10^. As one example, conventional resistance training exercises (i.e., calf muscle strengthening) have been shown to effectively increase not only muscle strength, but to also elicit tendon remodeling and increased stiffness^29^. As an alternative example, passive elastic ankle exoskeletons can augment the structural stiffness of ankle muscle-tendon units and their prescription may thus provide a benefit for metabolic cost in older adults^30^. The aforementioned interventions can directly augment Achilles tendon stiffness and thereby promote longer and more economical triceps surae muscle function. However, one notable challenge to the clinical translation of our findings is the direct vs. indirect consequences on reduced Achilles tendon stiffness with age on metabolic cost. Those with reduced k_AT_ may exhibit compensations at the hip which may play a larger role in increasing metabolic cost. Thus, more work in this area is warranted, to include precision joint-specific interventions – perhaps through the targeted use of assistive technologies.

There are several limitations of this work. One limitation is the lack of mechanistic data during walking to objectively link reduced k_AT_ with increased metabolic cost with more cause-effect precision. However, we have previously outlined in detail the neuromechanical cascade we believe is responsible for this association^17^. Another limitation is that we used EMG biofeedback to match activations between younger and older adults and not force. This was a purposeful methodological decision and should be used as motivation for further study. We also acknowledge that the significant correlation between reduced k_AT_ and increased walking metabolic cost was significant but relatively weak. This is not surprising, as the mechanisms governing increased walking metabolic cost due to age are likely numerous and complex. As one example, Hortobagyi et al. (2011) found that roughly one-third of the increase in metabolic cost could be explained by the economical consequences of antagonist muscle coactivation^31^. Finally, we acknowledge a relatively small sample size, particularly considering the presumably complex etiology of higher metabolic costs in the aging process and the diversity of age-related mobility impairments.

In conclusion, we provide empirical evidence that that lesser Achilles tendon stiffness in older than in younger adults exacts a metabolic penalty and is positively correlated with higher net metabolic power during walking. These results pave the way for interventions focused on restoring ankle muscle-tendon unit structural stiffness to improve walking energetics in aging.

## Acknowledgments

This study was supported by grants from the US National Institutes of Health (R01AG058615 and F32AG067675).

